## Supplementary information for "Regional response to light illuminance across the human hypothalamus"

[1]GIGA-Cyclotron Research Centre-In Vivo Imaging, University of Liège, Liège, 4000 Belgium
[2]Faculty of Health, Medicine and Life Sciences, School for Mental Health and Neuroscience, Alzheimer Centre Limburg, Maastricht University, Maastricht, 6229 ER The Netherlands.
[3]Institute of Neuroscience (IoNS), Department of Clinical Neuroscience,Université Catholique de Louvain (UCLouvain), Woluwe-Saint-Lambert, 1200 Belgium.
[4]Synergia Medical SA, Mont-Saint-Guibert, 1435 Belgium.
[#] shared first authorship
*Corresponding author: Gilles Vandewalle, GIGA-Cyclotron Research Centre-In Vivo Imaging, Bâtiment B30, 8 Allée du Six Août, University of Liège-Sart Tilman, 4000 Liège, Belgium.

**Supplementary Table S1. Demographics of study sample.**

| | Total Sample | Executive Task | Emotional Task |
|---|---|---|---|
| **Number of Participants** | 30 | 26 | 26 |
| **Age** | 24.2 ± 2.9 | 24.3 ± 3.0 | 24.4 ± 3.0 |
| **Sex (M)** | 11 | 10 | 10 |
| **Mood (BDI-II)** | 7.5 ± 7.0 | 6.7 ± 6.0 | 8.0 ± 7.3 |
| **Anxiety (BAI)** | 5.0 ± 4.1 | 4.8 ± 3.8 | 5.1 ± 4.3 |
| **Sleep quality (PSQI)** | 4.0 ± 2.6 | 3.7 ± 2.5 | 4.0 ± 2.7 |
| **Seasonality (SPAQ)** | 1.1 ± 0.8 | 1.2 ± 0.8 | 1.2 ± 0.8 |
| **Chronotype (HO)** | 48.7 ± 8.0 | 48.9 ± 8.2 | 48.7 ± 7.8 |
| **Daytime sleepiness (ESS)** | 6.5 ± 3.0 | 6.3 ± 3.0 | 6.2 ± 3.0 |
| **Years of Education** | 14.5 ± 3.1 | 14.5 ± 3.2 | 14.2 ± 3.2 |
| **Sleep duration (night before fMRI protocol – sleep diary based)** | 7.9 ± 0.7 | 7.8 ± 0.7 | 7.9 ± 0.7 |

Total number of participants who completed the study, and the number of participants included for each task (some participants had missing/corrupted data, see methods). BDI-II, Beck's Depression Inventory; BAI, Beck Anxiety Inventory; PSQI, Pittsburgh Sleep Quality Index; SPAQ, Seasonal Pattern Assessment Questionnaire; HO, Horne and Östberg; ESS, Epworth Sleepiness Scale. Refer to the method for the references to the scales and questionnaires.

**Supplementary Table S2. characteristics.**

| | Low BEL | Mid BEL | High BEL | Orange |
|---|---|---|---|---|
| **Lux** | 47 | 116 | 240 | 7.5 |
| **Peak Spectral Irradiance (nm)** | 460 | 460 | 460 | 590 |
| **Melanopic EDI (lux; ipRGCs)** | 37 | 92 | 190 | 0.16 |
| **Rhodopic EDI (lux; Rods)** | 39 | 97 | 201 | 0.94 |
| **Cyanopic EDI (lux; S-cones)** | 32 | 79 | 163 | 0 |
| **Chloropic EDI (lux; M-cones)** | 44 | 110 | 227 | 5 |
| **Erythropic EDI (lux ; L-cones)** | 46 | 113 | 233 | 8 |
| **Irradiance ($\mu W/cm^2$)** | 15 | 36 | 75 | 1.4 |
| **Photon flux($1/cm^2/s$)** | 4.12E+13 | 1.02E+14 | 2.10E+14 | 4.24E+12 |
| **Log Photon Flux ($log_{10}$ ($1/cm^2/s$)** | 13.61 | 14.01 | 14.32 | 12.63 |
| **Narrowband peak** | - | - | - | 589 |
| **Narrowband FWHM** | - | - | - | 10 |

Detailed characteristics of the four conditions used in fMRI protocol. Blue enriched (BEL) (low, mid, and high) and monochromatic (589nm). ipRGCs: intrinsically photosensitive retinal ganglion cells. FWHM: full width at half maximum.

**Supplementary Table S3. Post hoc contrasts between illuminances within each hypothalamus subpart during the executive task**

| Hypothalamus subpart | illuminance | vs. illuminance | t-value | p-value |
|---|---|---|---|---|
| **1 (inferior-anterior)** | 0 | 0.16 | 2.43 | **0.0151** |
| **1 (inferior-anterior)** | 0 | 37 | 2.59 | **0.0098** |
| **1 (inferior-anterior)** | 0 | 92 | 1.65 | 0.0993 |
| **1 (inferior-anterior)** | 0 | 190 | 3.30 | **0.0010** |
| **1 (inferior-anterior)** | 0.16 | 37 | 0.17 | 0.8683 |
| **1 (inferior-anterior)** | 0.16 | 92 | -0.78 | 0.4330 |
| **1 (inferior-anterior)** | 0.16 | 190 | 0.86 | 0.3886 |
| **1 (inferior-anterior)** | 37 | 92 | -0.95 | 0.3445 |
| **1 (inferior-anterior)** | 37 | 190 | 0.69 | 0.4892 |
| **1 (inferior-anterior)** | 92 | 190 | 1.65 | 0.0999 |
| **2 (superior-anterior)** | 0 | 0.16 | 0.79 | 0.4313 |
| **2 (superior-anterior)** | 0 | 37 | 0.76 | 0.4480 |
| **2 (superior-anterior)** | 0 | 92 | 1.37 | 0.1722 |
| **2 (superior-anterior)** | 0 | 190 | 1.15 | 0.2520 |
| **2 (superior-anterior)** | 0.16 | 37 | -0.02 | 0.9810 |
| **2 (superior-anterior)** | 0.16 | 92 | 0.58 | 0.5628 |
| **2 (superior-anterior)** | 0.16 | 190 | 0.36 | 0.7198 |
| **2 (superior-anterior)** | 37 | 92 | 0.60 | 0.5490 |
| **2 (superior-anterior)** | 37 | 190 | 0.38 | 0.7036 |
| **2 (superior-anterior)** | 92 | 190 | -0.22 | 0.8259 |
| **3 (posterior)** | 0 | 0.16 | -1.32 | 0.1873 |
| **3 (posterior)** | 0 | 37 | -1.22 | 0.2240 |
| **3 (posterior)** | 0 | 92 | -2.14 | **0.0323** |
| **3 (posterior)** | 0 | 190 | -2.35 | **0.0190** |
| **3 (posterior)** | 0.16 | 37 | 0.10 | 0.9180 |
| **3 (posterior)** | 0.16 | 92 | -0.82 | 0.4101 |
| **3 (posterior)** | 0.16 | 190 | -1.03 | 0.3037 |

| Hypothalamus subpart | illuminance | vs. illuminance | t-value | p-value |
|---|---|---|---|---|
| **3 (posterior)** | 37 | 92 | -0.93 | 0.3542 |
| **3 (posterior)** | 37 | 190 | -1.13 | 0.2578 |
| **3 (posterior)** | 92 | 190 | -0.21 | 0.8375 |
| **4 (inferior-tubular)** | 0 | 0.16 | 2.15 | **0.0316** |
| **4 (inferior-tubular)** | 0 | 37 | 2.21 | **0.0271** |
| **4 (inferior-tubular)** | 0 | 92 | 2.80 | **0.0052** |
| **4 (inferior-tubular)** | 0 | 190 | 3.27 | **0.0011** |
| **4 (inferior-tubular)** | 0.16 | 37 | 0.06 | 0.9518 |
| **4 (inferior-tubular)** | 0.16 | 92 | 0.65 | 0.5176 |
| **4 (inferior-tubular)** | 0.16 | 190 | 1.12 | 0.2624 |
| **4 (inferior-tubular)** | 37 | 92 | 0.59 | 0.5575 |
| **4 (inferior-tubular)** | 37 | 190 | 1.06 | 0.2891 |
| **4 (inferior-tubular)** | 92 | 190 | 0.47 | 0.6356 |
| **5 (superior-tubular)** | 0 | 0.16 | 0.01 | 0.9882 |
| **5 (superior-tubular)** | 0 | 37 | 0.84 | 0.3986 |
| **5 (superior-tubular)** | 0 | 92 | 0.86 | 0.3920 |
| **5 (superior-tubular)** | 0 | 190 | 0.58 | 0.5604 |
| **5 (superior-tubular)** | 0.16 | 37 | 0.83 | 0.4069 |
| **5 (superior-tubular)** | 0.16 | 92 | 0.84 | 0.4002 |
| **5 (superior-tubular)** | 0.16 | 190 | 0.57 | 0.5704 |
| **5 (superior-tubular)** | 37 | 92 | 0.01 | 0.9905 |
| **5 (superior-tubular)** | 37 | 190 | -0.26 | 0.7934 |
| **5 (superior-tubular)** | 92 | 190 | -0.27 | 0.7842 |

**Supplementary Table S4. Post hoc contrasts between illuminances within each hypothalamus subpart during the emotional task**

| Hypothalamus subpart | Illuminance | Vs. illuminance | t-value | p-value |
|---|---|---|---|---|
| **1 (inferior-anterior)** | 0 | 0.16 | -1.19 | 0.2324 |
| **1 (inferior-anterior)** | 0 | 37 | 1.29 | 0.1979 |
| **1 (inferior-anterior)** | 0 | 92 | 2.03 | **0.0431** |
| **1 (inferior-anterior)** | 0 | 190 | 2.25 | **0.0248** |
| **1 (inferior-anterior)** | 0.16 | 37 | 2.48 | **0.0132** |
| **1 (inferior-anterior)** | 0.16 | 92 | 3.22 | **0.0013** |
| **1 (inferior-anterior)** | 0.16 | 190 | 3.44 | **0.0006** |
| **1 (inferior-anterior)** | 37 | 92 | 0.74 | 0.4616 |
| **1 (inferior-anterior)** | 37 | 190 | 0.96 | 0.3379 |
| **1 (inferior-anterior)** | 92 | 190 | 0.22 | 0.8243 |
| **2 (superior-anterior)** | 0 | 0.16 | -0.14 | 0.8910 |
| **2 (superior-anterior)** | 0 | 37 | 1.14 | 0.2539 |
| **2 (superior-anterior)** | 0 | 92 | 2.86 | **0.0043** |
| **2 (superior-anterior)** | 0 | 190 | 3.49 | **0.0005** |
| **2 (superior-anterior)** | 0.16 | 37 | 1.28 | 0.2013 |
| **2 (superior-anterior)** | 0.16 | 92 | 3.00 | **0.0028** |
| **2 (superior-anterior)** | 0.16 | 190 | 3.63 | **0.0003** |
| **2 (superior-anterior)** | 37 | 92 | 1.72 | 0.0853 |
| **2 (superior-anterior)** | 37 | 190 | 2.35 | **0.0190** |
| **2 (superior-anterior)** | 92 | 190 | 0.63 | 0.5310 |
| **3 (posterior)** | 0 | 0.16 | -1.24 | 0.2151 |
| **3 (posterior)** | 0 | 37 | 0.13 | 0.8954 |
| **3 (posterior)** | 0 | 92 | -0.15 | 0.8799 |
| **3 (posterior)** | 0 | 190 | -2.17 | **0.0299** |
| **3 (posterior)** | 0.16 | 37 | 1.37 | 0.1704 |
| **3 (posterior)** | 0.16 | 92 | 1.09 | 0.2763 |
| **3 (posterior)** | 0.16 | 190 | -0.93 | 0.3506 |

| Hypothalamus subpart | Illuminance | Vs. illuminance | t-value | p-value |
|---|---|---|---|---|
| **3 (posterior)** | 37 | 92 | -0.28 | 0.7775 |
| **3 (posterior)** | 37 | 190 | -2.31 | **0.0213** |
| **3 (posterior)** | 92 | 190 | -2.02 | **0.0433** |
| **4 (inferior-tubular)** | 0 | 0.16 | 0.06 | 0.9486 |
| **4 (inferior-tubular)** | 0 | 37 | 1.01 | 0.3134 |
| **4 (inferior-tubular)** | 0 | 92 | 2.54 | **0.0113** |
| **4 (inferior-tubular)** | 0 | 190 | 2.42 | **0.0155** |
| **4 (inferior-tubular)** | 0.16 | 37 | 0.94 | 0.3454 |
| **4 (inferior-tubular)** | 0.16 | 92 | 2.47 | **0.0135** |
| **4 (inferior-tubular)** | 0.16 | 190 | 2.36 | **0.0185** |
| **4 (inferior-tubular)** | 37 | 92 | 1.53 | 0.1262 |
| **4 (inferior-tubular)** | 37 | 190 | 1.42 | 0.1571 |
| **4 (inferior-tubular)** | 92 | 190 | -0.11 | 0.9087 |
| **5 (superior-tubular)** | 0 | 0.16 | 0.04 | 0.9679 |
| **5 (superior-tubular)** | 0 | 37 | 1.85 | 0.0651 |
| **5 (superior-tubular)** | 0 | 92 | 1.71 | 0.0870 |
| **5 (superior-tubular)** | 0 | 190 | 1.10 | 0.2713 |
| **5 (superior-tubular)** | 0.16 | 37 | 1.81 | 0.0711 |
| **5 (superior-tubular)** | 0.16 | 92 | 1.67 | 0.0946 |
| **5 (superior-tubular)** | 0.16 | 190 | 1.06 | 0.2892 |
| **5 (superior-tubular)** | 37 | 92 | -0.13 | 0.8939 |
| **5 (superior-tubular)** | 37 | 190 | -0.75 | 0.4558 |
| **5 (superior-tubular)** | 92 | 190 | -0.61 | 0.5403 |

**Supplementary Table S5. Post hoc contrasts between hypothalamus subpart for each illuminance during the executive task**

| Illuminance | subpart | vs. subpart | t-value | p-value |
|---|---|---|---|---|
| **0** | 1 (inferior-anterior) | 2 (superior-anterior) | 1.25 | 0.2106 |
| **0** | 1 (inferior-anterior) | 3 (posterior) | 1.95 | 0.0511 |
| **0** | 1 (inferior-anterior) | 4 (inferior-tubular) | 0.37 | 0.7084 |
| **0** | 1 (inferior-anterior) | 5 (superior-tubular) | 0.84 | 0.4038 |
| **0** | 2 (superior-anterior) | 3 (posterior) | 0.70 | 0.4840 |
| **0** | 2 (superior-anterior) | 4 (inferior-tubular) | -0.88 | 0.3798 |
| **0** | 2 (superior-anterior) | 5 (superior-tubular) | -0.42 | 0.6763 |
| **0** | 3 (posterior) | 4 (inferior-tubular) | -1.58 | 0.1147 |
| **0** | 3 (posterior) | 5 (superior-tubular) | -1.12 | 0.2639 |
| **0** | 4 (inferior-tubular) | 5 (superior-tubular) | 0.46 | 0.6449 |
| **0.16** | 1 (inferior-anterior) | 2 (superior-anterior) | -0.13 | 0.8947 |
| **0.16** | 1 (inferior-anterior) | 3 (posterior) | -1.20 | 0.2287 |
| **0.16** | 1 (inferior-anterior) | 4 (inferior-tubular) | 0.14 | 0.8910 |
| **0.16** | 1 (inferior-anterior) | 5 (superior-tubular) | -1.20 | 0.2305 |
| **0.16** | 2 (superior-anterior) | 3 (posterior) | -1.07 | 0.2839 |
| **0.16** | 2 (superior-anterior) | 4 (inferior-tubular) | 0.27 | 0.7877 |
| **0.16** | 2 (superior-anterior) | 5 (superior-tubular) | -1.07 | 0.2860 |
| **0.16** | 3 (posterior) | 4 (inferior-tubular) | 1.34 | 0.1801 |
| **0.16** | 3 (posterior) | 5 (superior-tubular) | 0.00 | 0.9964 |
| **0.16** | 4 (inferior-tubular) | 5 (superior-tubular) | -1.34 | 0.1816 |
| **37** | 1 (inferior-anterior) | 2 (superior-anterior) | -0.29 | 0.7715 |
| **37** | 1 (inferior-anterior) | 3 (posterior) | -1.25 | 0.2105 |
| **37** | 1 (inferior-anterior) | 4 (inferior-tubular) | 0.05 | 0.9621 |
| **37** | 1 (inferior-anterior) | 5 (superior-tubular) | -0.64 | 0.5225 |
| **37** | 2 (superior-anterior) | 3 (posterior) | -0.96 | 0.3366 |
| **37** | 2 (superior-anterior) | 4 (inferior-tubular) | 0.34 | 0.7346 |
| **37** | 2 (superior-anterior) | 5 (superior-tubular) | -0.35 | 0.7279 |
| **37** | 3 (posterior) | 4 (inferior-tubular) | 1.31 | 0.1919 |

| Illuminance | subpart | vs. subpart | t-value | p-value |
|---|---|---|---|---|
| **37** | 3 (posterior) | 5 (superior-tubular) | 0.62 | 0.5382 |
| **37** | 4 (inferior-tubular) | 5 (superior-tubular) | -0.69 | 0.4904 |
| **92** | 1 (inferior-anterior) | 2 (superior-anterior) | 1.01 | 0.3107 |
| **92** | 1 (inferior-anterior) | 3 (posterior) | -1.24 | 0.2161 |
| **92** | 1 (inferior-anterior) | 4 (inferior-tubular) | 1.34 | 0.1801 |
| **92** | 1 (inferior-anterior) | 5 (superior-tubular) | 0.17 | 0.8668 |
| **92** | 2 (superior-anterior) | 3 (posterior) | -2.25 | **0.0246** |
| **92** | 2 (superior-anterior) | 4 (inferior-tubular) | 0.33 | 0.7438 |
| **92** | 2 (superior-anterior) | 5 (superior-tubular) | -0.85 | 0.3975 |
| **92** | 3 (posterior) | 4 (inferior-tubular) | 2.58 | **0.0101** |
| **92** | 3 (posterior) | 5 (superior-tubular) | 1.41 | 0.1602 |
| **92** | 4 (inferior-tubular) | 5 (superior-tubular) | -1.17 | 0.2409 |
| **190** | 1 (inferior-anterior) | 2 (superior-anterior) | -0.56 | 0.5782 |
| **190** | 1 (inferior-anterior) | 3 (posterior) | -2.80 | **0.0053** |
| **190** | 1 (inferior-anterior) | 4 (inferior-tubular) | 0.35 | 0.7229 |
| **190** | 1 (inferior-anterior) | 5 (superior-tubular) | -1.45 | 0.1479 |
| **190** | 2 (superior-anterior) | 3 (posterior) | -2.24 | **0.0254** |
| **190** | 2 (superior-anterior) | 4 (inferior-tubular) | 0.91 | 0.3626 |
| **190** | 2 (superior-anterior) | 5 (superior-tubular) | -0.89 | 0.3727 |
| **190** | 3 (posterior) | 4 (inferior-tubular) | 3.15 | **0.0017** |
| **190** | 3 (posterior) | 5 (superior-tubular) | 1.35 | 0.1781 |
| **190** | 4 (inferior-tubular) | 5 (superior-tubular) | -1.80 | 0.0718 |

**Supplementary Table S6. Post hoc contrasts <u>between</u> hypothalamus subpart for each illuminance during the <u>emotional task</u>**

| Illuminance | subpart | vs. subpart | t-value | p-value |
|---|---|---|---|---|
| **0** | 1 (inferior-anterior) | 2 (superior-anterior) | 0.45 | 0.6504 |
| **0** | 1 (inferior-anterior) | 3 (posterior) | 0.56 | 0.5775 |
| **0** | 1 (inferior-anterior) | 4 (inferior-tubular) | -0.34 | 0.7355 |
| **0** | 1 (inferior-anterior) | 5 (superior-tubular) | -1.50 | 0.1349 |
| **0** | 2 (superior-anterior) | 3 (posterior) | 0.10 | 0.9173 |
| **0** | 2 (superior-anterior) | 4 (inferior-tubular) | -0.79 | 0.4289 |
| **0** | 2 (superior-anterior) | 5 (superior-tubular) | -1.95 | 0.0515 |
| **0** | 3 (posterior) | 4 (inferior-tubular) | -0.90 | 0.3709 |
| **0** | 3 (posterior) | 5 (superior-tubular) | -2.05 | **0.0403** |
| **0** | 4 (inferior-tubular) | 5 (superior-tubular) | -1.16 | 0.2470 |
| **0.16** | 1 (inferior-anterior) | 2 (superior-anterior) | 1.38 | 0.1684 |
| **0.16** | 1 (inferior-anterior) | 3 (posterior) | 0.52 | 0.6049 |
| **0.16** | 1 (inferior-anterior) | 4 (inferior-tubular) | 0.76 | 0.4455 |
| **0.16** | 1 (inferior-anterior) | 5 (superior-tubular) | -0.42 | 0.6773 |
| **0.16** | 2 (superior-anterior) | 3 (posterior) | -0.86 | 0.3896 |
| **0.16** | 2 (superior-anterior) | 4 (inferior-tubular) | -0.62 | 0.5387 |
| **0.16** | 2 (superior-anterior) | 5 (superior-tubular) | -1.79 | 0.0730 |
| **0.16** | 3 (posterior) | 4 (inferior-tubular) | 0.25 | 0.8059 |
| **0.16** | 3 (posterior) | 5 (superior-tubular) | -0.93 | 0.3507 |
| **0.16** | 4 (inferior-tubular) | 5 (superior-tubular) | -1.18 | 0.2385 |
| **37** | 1 (inferior-anterior) | 2 (superior-anterior) | 0.32 | 0.7454 |
| **37** | 1 (inferior-anterior) | 3 (posterior) | -0.45 | 0.6497 |
| **37** | 1 (inferior-anterior) | 4 (inferior-tubular) | -0.58 | 0.5602 |
| **37** | 1 (inferior-anterior) | 5 (superior-tubular) | -1.01 | 0.3136 |
| **37** | 2 (superior-anterior) | 3 (posterior) | -0.78 | 0.4361 |
| **37** | 2 (superior-anterior) | 4 (inferior-tubular) | -0.91 | 0.3644 |
| **37** | 2 (superior-anterior) | 5 (superior-tubular) | -1.33 | 0.1828 |
| **37** | 3 (posterior) | 4 (inferior-tubular) | -0.13 | 0.8979 |

| Illuminance | subpart | vs. subpart | t-value | p-value |
|---|---|---|---|---|
| **37** | 3 (posterior) | 5 (superior-tubular) | -0.55 | 0.5798 |
| **37** | 4 (inferior-tubular) | 5 (superior-tubular) | -0.43 | 0.6706 |
| **92** | 1 (inferior-anterior) | 2 (superior-anterior) | 1.19 | 0.2355 |
| **92** | 1 (inferior-anterior) | 3 (posterior) | -1.35 | 0.1788 |
| **92** | 1 (inferior-anterior) | 4 (inferior-tubular) | 0.11 | 0.9111 |
| **92** | 1 (inferior-anterior) | 5 (superior-tubular) | -1.77 | 0.0772 |
| **92** | 2 (superior-anterior) | 3 (posterior) | -2.53 | **0.0115** |
| **92** | 2 (superior-anterior) | 4 (inferior-tubular) | -1.08 | 0.2825 |
| **92** | 2 (superior-anterior) | 5 (superior-tubular) | -2.96 | **0.0032** |
| **92** | 3 (posterior) | 4 (inferior-tubular) | 1.46 | 0.1454 |
| **92** | 3 (posterior) | 5 (superior-tubular) | -0.42 | 0.6721 |
| **92** | 4 (inferior-tubular) | 5 (superior-tubular) | -1.88 | 0.0603 |
| **190** | 1 (inferior-anterior) | 2 (superior-anterior) | 1.54 | 0.1237 |
| **190** | 1 (inferior-anterior) | 3 (posterior) | -3.31 | **0.0010** |
| **190** | 1 (inferior-anterior) | 4 (inferior-tubular) | -0.18 | 0.8549 |
| **190** | 1 (inferior-anterior) | 5 (superior-tubular) | -2.50 | **0.0126** |
| **190** | 2 (superior-anterior) | 3 (posterior) | -4.85 | **<.0001** |
| **190** | 2 (superior-anterior) | 4 (inferior-tubular) | -1.72 | 0.0851 |
| **190** | 2 (superior-anterior) | 5 (superior-tubular) | -4.04 | **<.0001** |
| **190** | 3 (posterior) | 4 (inferior-tubular) | 3.13 | **0.0018** |
| **190** | 3 (posterior) | 5 (superior-tubular) | 0.81 | 0.4182 |
| **190** | 4 (inferior-tubular) | 5 (superior-tubular) | -2.32 | **0.0208** |

**Supplementary Table S7. Association between performance to the 2-back task and the activity of each hypothalamus subpart during each illuminance**

| | F-value | p-value | Partial $R^2$ |
|---|---|---|---|
| **1 (inferior-anterior hypothalamus subpart)** | | | |
| **Subpart activity** | < 0.01 | 0.99 | |
| **Illuminance** | 1.94 | 0.13 | |
| **Age** | 0.04 | 0.84 | |
| **Sex** | 6.43 | **0.019** | 0.23 |
| **BMI** | 2.02 | 0.16 | |
| **2 (superior-anterior hypothalamus subpart)** | | | |
| **Subpart activity** | 0.62 | 0.43 | |
| **Illuminance** | 2.24 | 0.07 | |
| **Age** | 0.01 | 0.94 | |
| **Sex** | 6.36 | **0.019** | 0.22 |
| **BMI** | 2.04 | 0.17 | |
| **3 (Posterior hypothalamus subpart)** | | | |
| **Subpart activity** | 9.43 | **0.0027** | 0.08 |
| **Illuminance** | 2.72 | **0.034** | 0.1 |
| **Age** | 0.04 | 0.85 | |
| **Sex** | 6.07 | **0.022** | 0.21 |
| **BMI** | 1.82 | 0.19 | |
| **4 (inferior-tubular hypothalamus subpart)** | | | |
| **Subpart activity** | 0.12 | 0.7 | |
| **Illuminance** | 2.09 | 0.11 | |
| **Age** | 0.03 | 0.86 | |
| **Sex** | 6.54 | **0.018** | 0.23 |
| **BMI** | 2.01 | 0.17 | |
| **5 (superior-tubular hypothalamus subpart)** | | | |
| **Subpart activity** | 0.25 | 0.62 | |
| **Illuminance** | 2.12 | 0.084 | |
| **Age** | 0.02 | 0.88 | |
| **Sex** | 6.1 | **0.021** | 0.21 |
| **BMI** | 2.01 | 0.17 | |
